# Constitutive Androstane Receptor and Hepatitis B Virus X Protein Cooperatively Induce β-catenin-Activated Liver Tumors

**DOI:** 10.1101/2020.08.08.241661

**Authors:** Jessica D. Scott, Silvia Liu, Kevin C. Klatt, Zhen Sun, Qi Guo, Sandra L. Grimm, Cristian Coarfa, Bingning Dong, David D. Moore

**Affiliations:** Department of Molecular and Cellular Biology, Baylor College of Medicine, Houston, TX; Integrative Molecular and Biomedical Sciences Graduate Program, Baylor College of Medicine, Houston, TX; Department of Pathology, School of Medicine, University of Pittsburgh, Pittsburgh, PA

## Abstract

**Background and Aims:** The xenobiotic nuclear receptor Constitutive Androstane Receptor (CAR) is essential for xenobiotic tumor promotion in mouse models. In these models, β-catenin is genetically activated in approximately 80% of tumors. Chronic Hepatitis B Virus (HBV) infection is a major risk factor for hepatocellular carcinoma (HCC), and β-catenin activation is also frequently activated in HBV-associated HCCs. The goal of this research was to determine whether activation of CAR in a mouse model of chronic HBV infection would result in tumor formation and whether these tumors would display increased β-catenin activation.

**Approach and Results:** We treated transgenic mice expressing the HBV X protein (HBx) in hepatocytes with a single dose of the potent CAR agonist TCPOBOP. After 10 months, these mice developed large liver tumors that are characterized by β-catenin nuclear localization and upregulation of β-catenin targets. The β-catenin regulator FoxM1 and the oxidative stress master regulator Nrf2, both of which are CAR gene targets, were also overactivated in tumors. The CAR/HBx tumors share a conserved gene signature with HBV-related human hepatocellular carcinoma.

**Conclusions:** Activation of CAR in the presence of HBx results in tumors with strong β-catenin activation. The mouse model we have described reflects the gene expression patterns seen in human HBV-associated HCC and presents an attractive basis for future studies.

---

Hepatocellular carcinoma (HCC) is the most common form of liver cancer, affecting over 14 million people worldwide (1,2). It is one of the fastest-growing causes of death in the United States (1). Despite improved screening and diagnostics, HCC incidence continues to rise, and 5-year survival averaged over all stages remains only 4-12% (3–5).

The biggest risk factors for developing HCC include chronic Hepatitis B (HBV) or Hepatitis C (HCV) infection, alcoholic liver cirrhosis, and nonalcoholic fatty liver disease (NAFLD) (1). Of these, chronic HBV infection is the single greatest predictor for developing HCC worldwide. Approximately 15-25% of chronic HBV sufferers will die from cirrhosis or HCC (6). Despite widespread adoption of the HBV vaccine in the 1990s and 2000s, HCC due to chronic HBV infection continues to rise, in part because chronic HBV infection is often contracted in early infancy (7).

In studies examining hierarchical clustering of HCC tumors, HBV-associated tumors do not form a distinct cluster (8,9), indicating that analysis of these tumors may yield insights that are applicable across many HCC etiologies. HBV X protein (HBx) is the sole regulatory protein encoded in the compact genome of HBV and plays important roles in viral propagation, gene transcription, and alteration of host signaling pathways. It is also thought to contribute to HCC development by affecting cell signaling, apoptosis, DNA repair, and cell cycle progression in hepatocytes (10).

The nuclear receptor Constitutive Androstane Receptor (CAR, NR1I3) was first characterized as an important regulator of xenobiotic metabolism in the liver (11). More recently, CAR has also been implicated in cell proliferation and liver tumor development. Acute CAR activation causes hepatocyte replication, resulting in limited, reversible liver growth (12). Chronic CAR activation is sufficient to induce limited numbers of tumors in rodent models. However, when combined with pretreatment with a genotoxic tumor initiator such as diethylnitrosamine (DEN), CAR activation strongly promotes hepatocarcinogenesis (13). CAR is essential for tumor promotion by phenobarbital, an indirect CAR activator, and by TCPOBOP, a direct CAR agonist (12,13). The oncogenic response to CAR activation appears to be linked to β-catenin. In phenobarbital-induced liver tumors, a remarkable 80% contain activating mutations in *CTNNB1*, the gene that encodes β-catenin (14). We previously reported that the acute pharmacologic activation of CAR combined with genetic overactivation of β-catenin caused extensive liver proliferation and tumorigenesis (15). These CAR/β-catenin tumors share a conserved gene signature with the subset of human HCC that have activating β-catenin mutations (15).

Wnt/β-catenin signaling is active in liver development as well as during liver regeneration following resection or toxicity. Activated β-catenin translocates from the cytoplasm to the nucleus, where it induces target genes that are involved in cell cycle progression, survival, and differentiation (16). Dysregulation of the Wnt/β-catenin pathway is a common event in many types of cancer, including HCC. Approximately 30-50% of human HCC tumors contain activating mutations in the Wnt/β-catenin pathway or nuclear accumulation of β-catenin (17–19). Alterations to the Wnt/β-catenin pathway are common across HCC tumors, regardless of etiology. Mutations to *CTNNB1* itself are more common in HCV-related tumors, whereas HBV-related carcinomas are more likely to contain alterations to genes encoding β-catenin regulatory proteins such as GSK-3β or AXIN1/2 (20). Notably, HBx has been reported to induce β-catenin activation; however, mice expressing an HBx transgene in hepatocytes typically do not develop spontaneous liver tumors and require additional factors to induce tumorigenesis (21,22). We previously reported that CAR synergizes with β-catenin to induce liver tumors (15). This interaction suggests that CAR may also play a role in HBV-mediated progression of HCC.

We report here that CAR and HBx functionally interact to induce tumorigenesis via β-catenin activation. We found that a single injection of the potent CAR agonist TCPOBOP in 8-week-old HBx transgenic mice (ATX mice) consistently results in liver tumors by 12 months of age. Strong β-catenin activation is a hallmark of these tumors. The FoxM1 and Nrf2 transcription factor pathways, which promote HCC progression and are linked to poor prognosis in human HCC (23,24), were also upregulated in the CAR/HBx tumors. Interestingly, while 8-week-old ATX mice have mildly increased β-catenin target expression compared to wildtypes, acute CAR activation appears to suppress β-catenin activation. Addition of the CAR knockout allele to the HBx transgene completely prevented tumor development and induction of gene and protein levels of tumor-associated targets. We propose that CAR functionally interacts with HBx to promote β-catenin-mediated liver tumorigenesis.

## Materials and Methods

### Animal Maintenance and Treatment

All mice used in these experiments were 8-10 weeks old at the start of treatment. 3 mg/kg TCPOBOP or equivalent volume of Corn Oil was administered to mice via a single intraperitoneal injection. Immediately prior to tissue collection at the indicated time points, mice were deeply anesthetized by isofluorane inhalation and then euthanized by cervical dislocation. All experiments involving mice were approved by the Institutional Animal Care and Use Committee of Baylor College of Medicine. All mice were housed in the Baylor College of Medicine Transgenic Mouse Facility.

### ATX Mouse Generation

ATX breeding pairs in the ICR background were generously provided by the Slagle laboratory. The genetics of ATX mice have been previously described (21). Breeding pairs included one ATX-positive mouse and one ATX-negative mouse. Wildtype and ATX mice used in experiments came from the same litters. All ATX mice were hemizygous for the transgene. Presence or absence of transgene was confirmed by genotyping.

### Generation of CAR -/- Mice

ATX-positive mice in the ICR background were bred to CAR -/- mice. Heterozygous CAR, ATX-negative mice from this mating were then back-crossed to ATX-positive ICR mice. In order to ensure complete integration of the ICR background, CAR -/- mice were subsequently back-crossed to ICR mice for at least nine additional generations before experiments were performed.

### Liver Histology

Fresh liver samples measuring approximately 1 cm^2^ were removed and fixed in 10% Neutral Buffered Formalin. Samples were embedded in paraffin and sectioned into 5 um slices onto charged slides. Slides were either stained with Hematoxylin and Eosin or underwent antigen retrieval and antibody staining. Antigen retrieval was performed using heat-mediated Sodium Citrate Buffer. Tissues were permeabilized with 0.1% Triton X-100. Sections were stained with β-catenin antibody (1:400), GS antibody (1:400), AFP (1:200), Ki-67 (1:400), or FoxM1 (1:400). Ki67 quantification was performed by counting a minimum of 200 cells from at least four randomly selected fields per tissue sample. At least three samples were included per group.

### RNA Isolation and Real-Time Quantitative PCR

Mouse tissues were collected and rapidly frozen in liquid nitrogen. Total RNA was isolated using TRIzol reagent (GenDEPOT). Tissue was homogenized in two 10-second intervals using MagNA Lyser (Roche), incubated with 200 ul chloroform (Fisher Scientific) with mixing for two minutes, and centrifuged for 12,000xg at 4°C for 15 minutes. RNA was collected from the supernatant and cleaned using the Quick-RNA MiniPrep Kit (Zymo). RNA was converted to cDNA using the qScript cDNA Synthesis Kit (Quantabio, 95047). Cycle threshold values were obtained by Sybr Green (KAPA) detection using the LightCycler 480 System (Roche). Relative expression was quantified by delta delta Ct analysis normalized to Cyclophilin A or GAPDH. Primer sequences may be found in Table S1.

### Protein Isolation and Western Blotting

Mouse tissues were collected and rapidly frozen in liquid nitrogen. 5-10 mg of tissue was sonicated in 500 ul Radioimmunoprecipitation Assay (RIPA) buffer supplemented with Protease Inhibitor (GenDEPOT) and Phosphatase Inhibitor (GenDEPOT). Lysate was centrifuged at 14,000 x g and total protein was collected from the supernatant. Protein concentration was measured by BCA assay (Thermo Scientific), which was read at 550 nm (Multiskan FC, Thermo Scientific). 30-50 ug of protein was loaded onto a 4-15% gel (Bio-Rad), run in Tris-Glycine buffer at 100V for 2 hours, and transferred at 20V overnight onto a nitrocellulose membrane (Immobilon P^SQ^, Millipore) using an all-wet transfer system (Bio-Rad). Membranes were blocked for 1 hour at room temperature with 5% milk protein. Antibodies were incubated at 4° C overnight in 5% milk protein at an antibody-dependent dilution. Antibodies used included Glutamine Synthetase (BD Biosciences; 1:2000) and β-actin (Cell Signaling; 1:4000).

### RNA-Sequencing

Mouse tissues were collected and rapidly frozen in liquid nitrogen. 5-10 mg of tissue was placed in a tube containing ceramic beads along with 500 ul PureXtract RNAsol reagent (GenDEPOT). RNA was collected and cleaned as described above. RNA quality was verified using Qubit 3.0 fluorometer and Agilent TapeStation System. The cDNA library was prepared using the TruSeq Stranded mRNA Sample Preparation Kit (Illumina). IDT TruSeq RNA UD Indexes (Illumina) were used for the adapter index. Paired-end sequencing was performed on the NextSeq 550 System (Illumina) using the NextSeq 500/550 v2.5 Kit (Illumina). FPKM values for 11730 genes across 4 wild type (WC) and 5 HBV-induced tumor samples (AT) were collected. Test for differential expression was performed by R package ‘limma’. Differentially expressed genes (DEGs) were defined by FDR=5% and absolute fold change greater than 2. These genes were further used for input into Ingenuity Pathway Analysis (IPA) ^®^. Significant pathways were selected by FDR=5%.

### Statistical Analysis

Comparison of gene expression between tissue groups was analyzed by one-way analysis of variance. Student’s t-test was applied to obtain p-values between two groups. Statistical significance was defined as p < 0.05. Data are represented as mean ± SEM. All statistical tests were performed using GraphPad Prism.

### Human data comparison

Three public human HCC studies were explored in this study: GSE121248, GSE94660 and TCGA LIHC project. For GSE121248 study, gene expression for 37 adjacent normal samples and 70 HBV-infected HCC tumors were collected. Microarray probe intensities were mapped to gene expressions. If multiple probes were mapped to the same gene, the probe with the largest interquartile range was used as representative of that gene. Five outlier samples were removed based on principal component analysis. Eventually 22880 genes across 36 adjacent normal samples and 66 HBV-infected HCC tumors were used for the downstream analysis. For GSE94660 study, FPKM values for 26760 genes across 21 paired non-tumor and HBV-infected tumors samples were analyzed. For TCGA LIHC project, FPKM values of 58387 genes across 50 normal sample, 101 HBV-infected tumors and 268 non-HBV-infected tumors were collected for analysis. For each of these three studies, differential expression analysis and pathway analysis were performed with the same pipeline as mouse model.

### SNVs calling on RNA-seq data

Raw RNA-sequencing reads were first trimmed by tool Trimmomatic to filter out low-quality reads and adapter sequences. Then survived reads were aligned to mouse reference genome mm10 by STAR aligner. Based on the aligned reads, SNVs/mutations were called on each individual sample by SAMtools and bcftools mpileup function. Known SNPs and genes were annotated by SnpEff and SnpSift toolbox.

## Results

### Activation of CAR in mice expressing HBx leads to tumorigenesis

CAR is essential for liver tumor promotion in mice, but it has a limited ability to induce tumors on its own. Pretreatment with strong tumor initiators such as the genotoxic carcinogen diethylnitrosamine (DEN) is used in conjunction with CAR activation to induce liver tumors. These models, while useful, do not imitate physiologically relevant conditions for natural tumor formation. In order to partially mimic consequences of chronic HBV infection, we used HBx transgenic mice (ATX mice), which have been previously characterized (21,22). Presence of the ATX transgene was confirmed by genotyping and by RT-qPCR (Supplemental Figure S1). Primer sequence is given in Table S1. Wildtype littermates were used as controls. CAR was activated in eight-week-old mice with a single injection of TCPOBOP, a potent and persistent CAR agonist, and assessed at one year of age. Ten months after treatment with TCPOBOP, ATX mice developed large tumors throughout the liver (Figure 1A). All ATX mice treated with TCPOBOP incurred liver tumors, compared with one-third of wildtype TCPOBOP-treated mice (Figure 1B). In general, tumors in wildtype mice, when present, were smaller and less advanced than those occurring in ATX mice. Vehicle-treated wildtype mice were completely tumor-free, and only one very small tumor nodule was observed in vehicle-treated ATX mice. In order to confirm that tumors were CAR-dependent, we repeated the experiment in CAR null mice. No tumors were observed in CAR null mice without the HBx transgene (Figure 1A).

**Figure 1.**
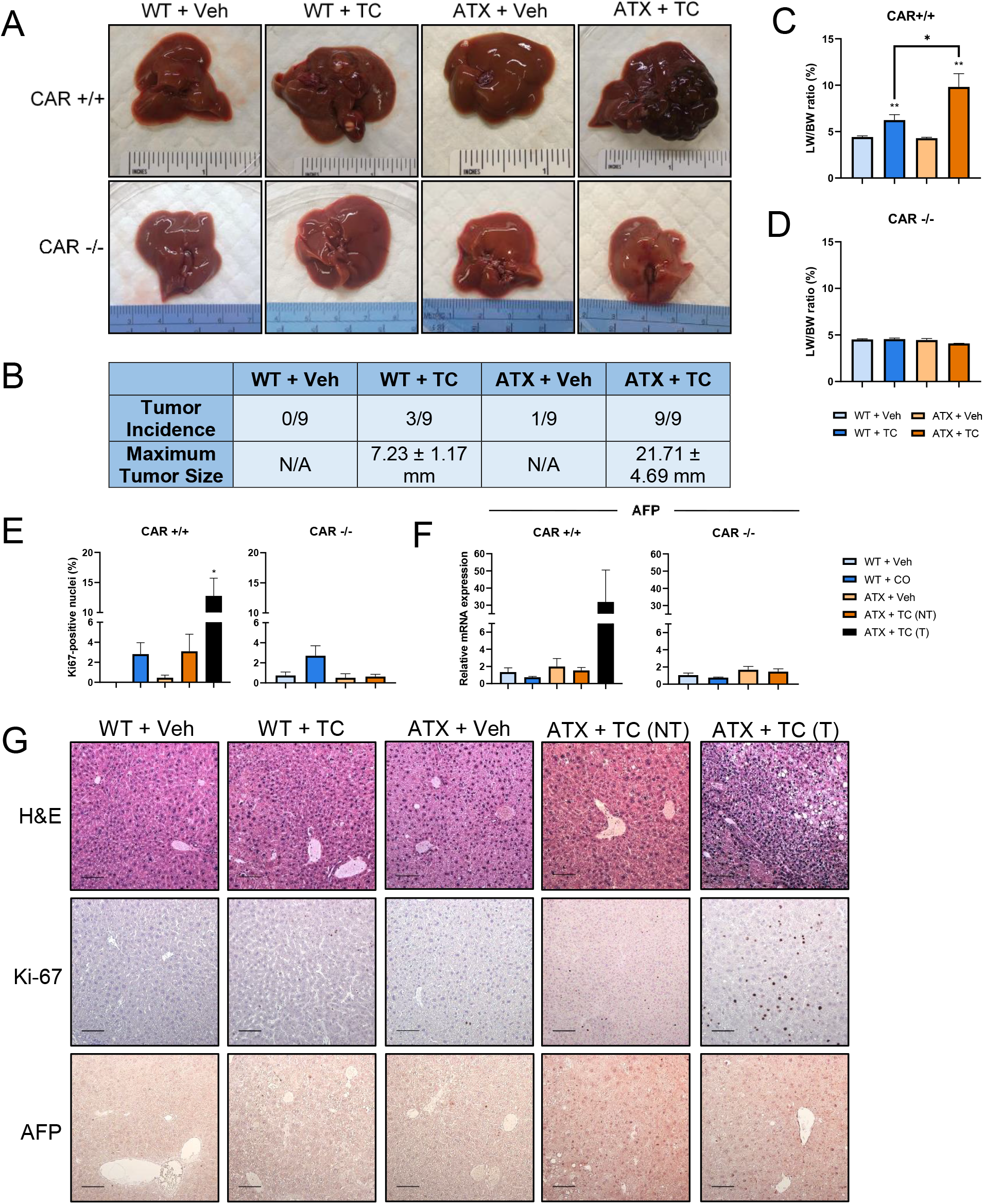
Long-term CAR activation in ATX mice leads to liver tumorigenesis. Wild-type or ATX mice were injected with a single dose of 3 mg/kg TCPOBOP or vehicle at 8 weeks of age and analyzed at 12 months of age. (A) Representative gross appearances of livers. (B) Tumor incidence and maximum tumor size in each group. (C) Liver-to-body weight ratios. (D) Percent of Ki-67-positive cells. (E) Relative RNA expression of AFP as measured by qPCR. * p < 0.05 (F) Immunohistochemistry of Ki-67 and alpha-fetoprotein (AFP) in liver sections. Scale bars represent 100 μm.

As expected, TCPOBOP-treated wildtype mice had a modest but significant increase in liver-to-body weight ratio. Liver-to-body weight ratios of vehicle-treated ATX mice were not different from wildtype, whereas liver-to-body weight ratio of TCPOBOP-treated ATX mice more than doubled their liver-to-body weight ratio (Figure 1C). CAR null mice had no difference in liver-to-body weight ratio regardless of treatment (Figure 1D). The percentage of Ki67-positive nuclei was significantly elevated in ATX + TCPOBOP tumors. There was a slight, but not statistically significant, increase in Ki67-positive nuclei in both wildtype + TCPOBOP and ATX + TCPOBOP nontumor tissues, and no significant elevation of Ki67 in CAR null tissues (Figure 1 E, G). Alpha fetoprotein (AFP) is normally expressed in developing fetal liver tissue and is commonly used as a marker of hepatocellular carcinoma. AFP gene expression was highly upregulated in tumors and remained low in all other groups (Figure 1 F-G). Histology of CAR null mice, as well as AFP and Ki-67 levels, were normal (Supplemental Figure S2).

### β-catenin activation in CAR/ATX tumors

HBx is reported to activate β-catenin, though this effect is not directly transforming (25,26). Because of the known synergistic effect of CAR and β-catenin, we tested whether tumors induced by CAR activation in ATX mice were associated with β-catenin activation. In ATX + TCPOBOP tumors, we found β-catenin to be strongly upregulated. This effect was not observed in other treatment groups, nor in non-tumor tissue. We saw a clear pattern of β-catenin nuclear localization in tumor tissues, as well as widespread expression of glutamine synthetase, a direct β-catenin target (Figure 2A). In all other groups, glutamine synthetase follows a typical zonal pattern of expression, only surrounding hepatic veins. Glutamine synthetase in tumor tissues is broadly and strongly expressed throughout the tumor. In addition, we found that gene expression of β-catenin targets, including glutamine synthetase, GPR49, Slc1a2, and TCF, was upregulated in tumors (Figure 2C). Expression of FoxO1, which is known to suppress β-catenin activity (27), was significantly reduced in both tumor and non-tumor tissues (Figure 2C). We also found that protein expression of glutamine synthetase was highly upregulated in tumor tissue compared to other tissues (Figure 2A, D). CAR knockout ablated the β-catenin activation phenomenon observed in CAR +/+ tumors. We did not observe any β-catenin nuclear localization for any CAR null groups (Figure 2B). Additionally, glutamine synthetase maintained its normal, peri-venous protein expression pattern. mRNA expression of β-catenin targets did not differ between groups (data not shown).

**Figure 2.**
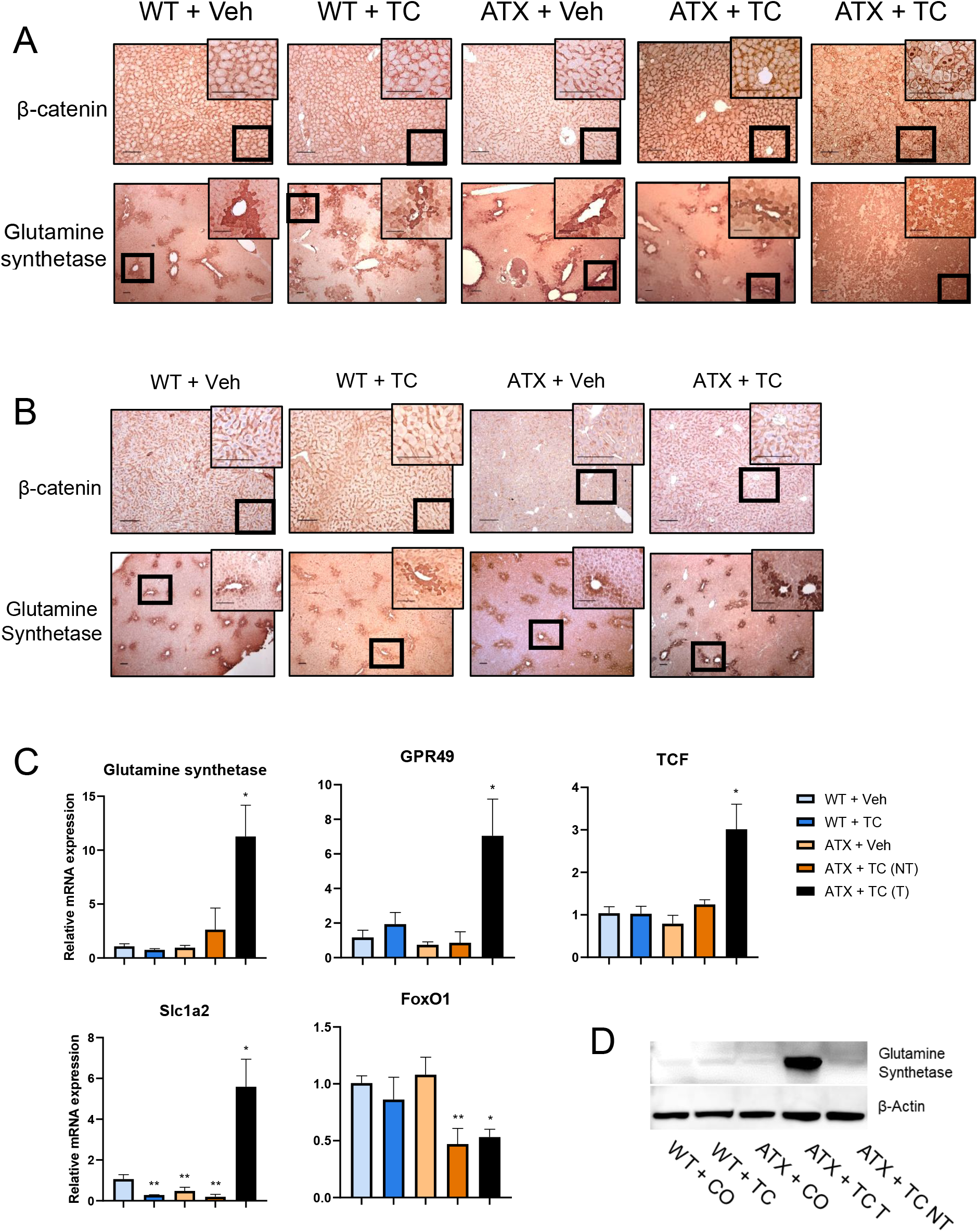
CAR activation in ATX mice causes β-catenin activation. (A) Hematoxylin and eosin staining; β-catenin and glutamine synthetase staining. (B) qPCR analysis of β-catenin-related gene expression. (C) qPCR analysis of FoxM1-related gene expression. * p < 0.05; ** p < 0.01. (D) Representative Western blot of glutamine synthetase.

### The FoxM1 and Nrf2 pathways are activated in ATX + TC tissues

The pro-proliferative transcription factor FoxM1 is known to promote β-catenin nuclear localization (28). In a previous study, we found that dual activation of CAR and β-catenin upregulates FoxM1 pathway genes (15). We therefore tested whether FoxM1 is activated in ATX + TCPOBOP-treated tumor tissues. Increased nuclear localization is evident in both tumor and non-tumor ATX + TCPOBOP tissues (Figure 3A). FoxM1 gene expression and target gene expression were also increased (Figure 3B).

**Figure 3.**
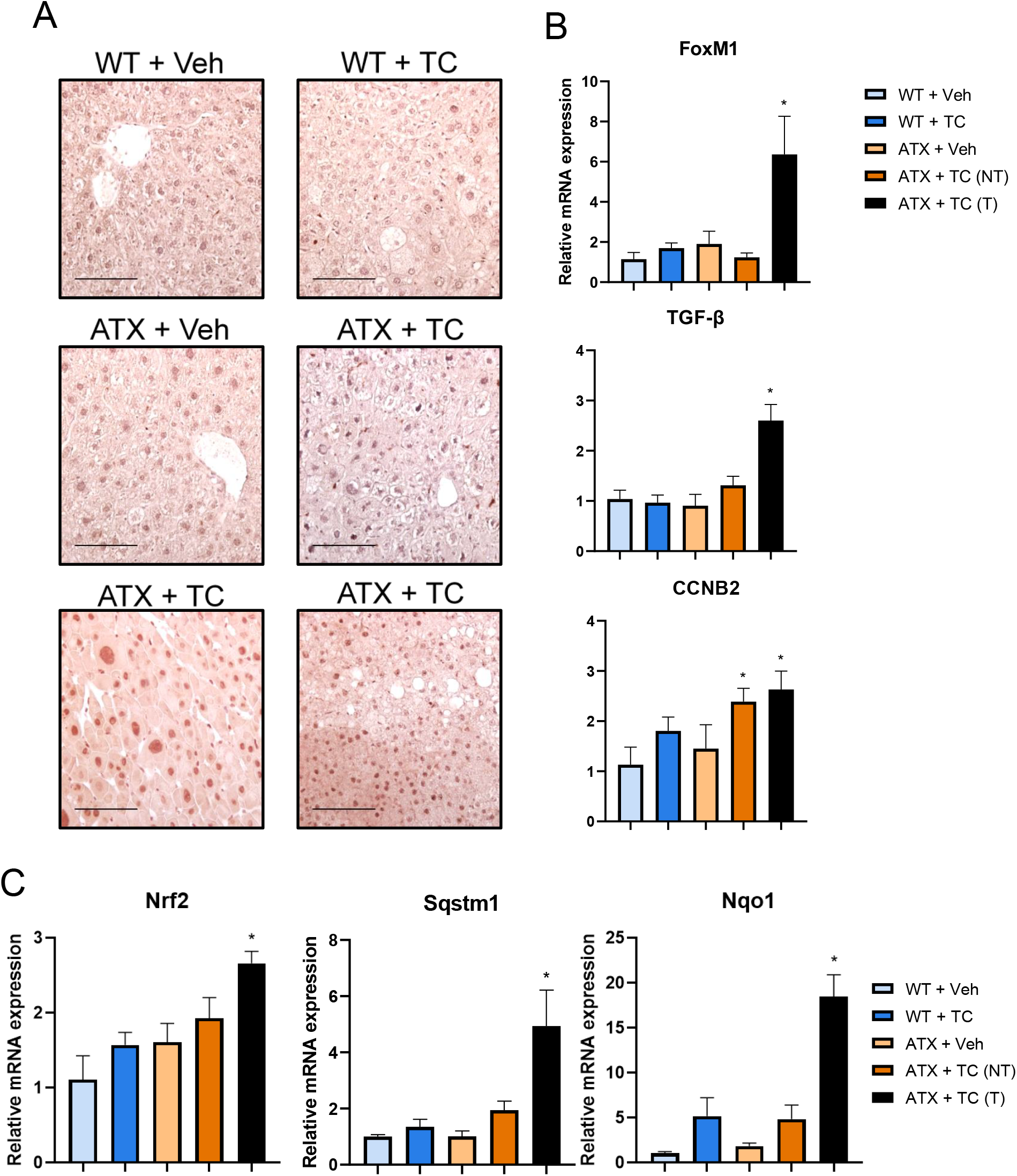
FoxM1 is activated in ATX + TC tissues. (A) FoxM1 tissue staining. B) qPCR analysis of FoxM1-related gene expression. * p < 0.05.

FoxM1 regulates expression of the antioxidant response transcription factor Nrf2 (29,30). Nrf2 promotes cell survival and proliferation in response to oxidative stress conditions, often an environment inherent in HCC development (23). Nrf2 activation is tightly linked to CAR activation, likely through CAR-mediated induction of oxidative stress (31). Additionally, Nrf2 activation can be stimulated by HBV infection, specifically by HBx activity. We therefore suspected that Nrf2 pathways would be upregulated in our tumor model. Indeed, we observed increased gene expression of Nrf2 and its target genes Nqo1 and Sqstm1 in tumor tissues (Figure 3C). Increases in Nrf2 activity have been found in tissue samples from HBV-positive HCC samples, making this a potential therapeutic target for this patient population (32).

### Short-term CAR activation in ATX mice leads to hepatocyte proliferation

In order to assess changes that occur at early timepoints, we treated mice as above and sacrificed three days later. As expected, livers treated with TCPOBOP were larger than livers of vehicle-treated mice (Figure 4A). Both groups treated with TCPOBOP had significantly higher liver-to-body weight ratios than control groups, with ATX + TCPOBOP differing significantly from the wildtype TCPOBOP-treated group (Figure 4B). Hematoxylin and eosin staining of these tissues can be found in Figure 4C. TCPOBOP produced an extremely strong proliferative effect in both genotypes, as evidenced by extensive Ki-67 staining (Figure 4B, C). We did not observe a significant difference in Ki-67 staining between the two TCPOBOP-treated groups.

**Figure 4.**
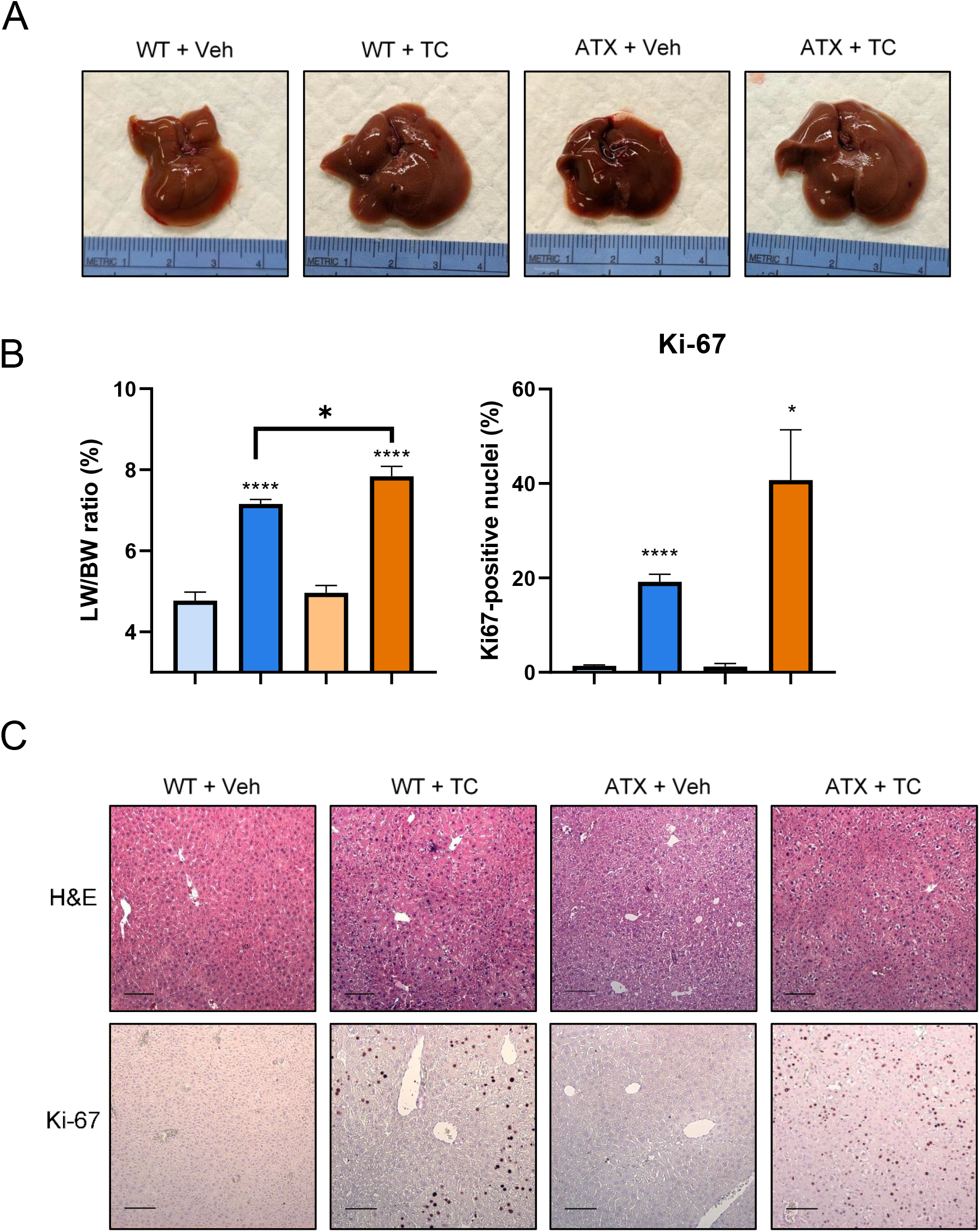
Short-term CAR activation in ATX mice causes hepatocyte proliferation. Wild-type or ATX mice were injected with a single dose of 3 mg/kg TCPOBOP or vehicle at 8 weeks of age and analyzed at 12 months of age. (A) Representative gross appearances of livers. (B) Liver-to-body weight ratios; percent of positive Ki-67 cells. Groups were analyzed using two-way analysis of variance. (C) Hematoxylin and eosin staining; immunohistochemistry of Ki-67 and FoxM1. Scale bars represent 100 μm.

### Induction of the FoxM1 and Nrf2 pathways, but not β-catenin activation, are early events in CAR-mediated transformation of ATX livers

In order to identify early events that may contribute to increased hepatocyte proliferation, we assessed mice in the groups described above at three days post-treatment. No β-catenin nuclear localization was apparent by immunohistochemistry (Figure 5C). Glutamine synthetase was also expressed normally in all tissue types. Because HBx is reported to induce β-catenin pathways, we expected to see an increase in β-catenin target genes in 8-week-old mice. While we did see slight increases in target gene expression, most of these changes were not statistically significant. Based on the functional synergy apparent from the tumors and the previously described acute responses (15), we anticipated that CAR activation would increase β-catenin responses. Unexpectedly, however, acute CAR activation lowered β-catenin target gene expression. In all cases, TCPOBOP-treated ATX tissues expressed β-catenin target genes at a significantly lower level than vehicle-treated ATX tissues (Figure 5A).

**Figure 5.**
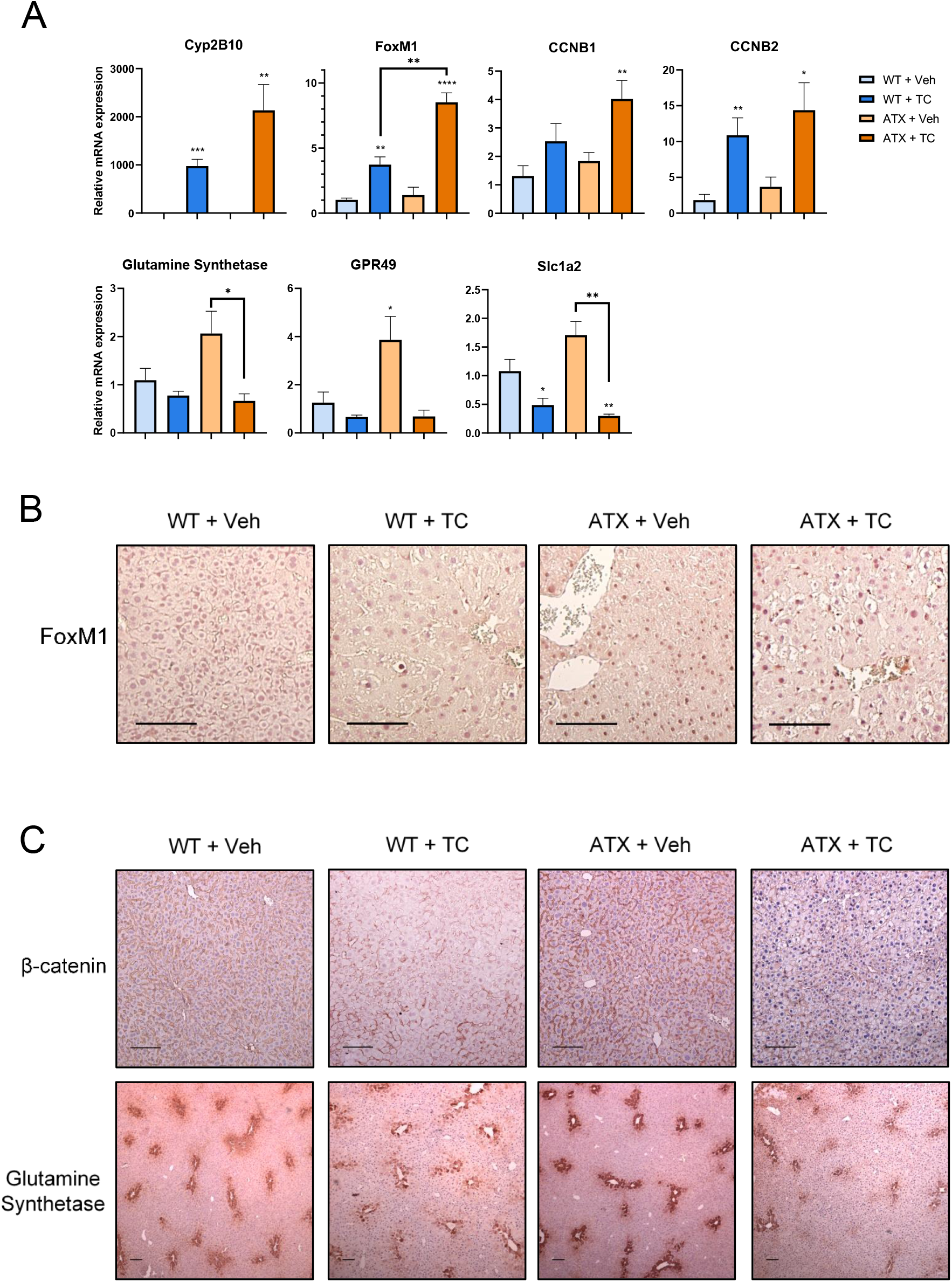
FoxM1 and Nrf2 upregulation are early events in CAR-activated ATX mice. (A) qPCR analysis of gene expression. * p < 0.05; ** p < 0.01; *** p < 0.001; **** p < 0.0001. (B) Immunohistochemistry of FoxM1. (C) Immunohistochemistry of β-catenin and glutamine synthetase.

We hypothesized that activation of FoxM1, a previously described CAR target that is involved in liver hyperplasia and cancer (24,33), would be an early event in CAR-mediated transformation of ATX liver. Indeed, we discovered that TCPOBOP treatment of ATX mice results in upregulation of mRNA levels of FoxM1 as well as its targets (Figure 5A). These findings are corroborated by immunohistochemistry of FoxM1, which displays increased nuclear staining in ATX + TCPOBOP tissues (Figure 5B).

Since previous results identify Nrf2 as a direct target of both CAR and FoxM1 (29,31), we hypothesized that Nrf2 would also constitute an early event in ATX + TCPOBOP-induced proliferation. As expected, we observed significant increases in Nrf2 targets at early timepoints (Figure 5A).

### Transcriptome analysis on mouse model

In order to study the transcriptome profiles of the mouse model, RNA sequencing was performed on 4 wild type liver (WC) samples from non-tumor bearing mice and 5 CAR/HBV-induced tumor samples (AT). Principal component analysis (PCA) was performed based on the FPKM (fragments per kilobase per million reads) values. As expected, wild-type samples (blue dots) and tumor samples (orange triangles) clustered in two separate groups on Principal Component analysis. Differential expression analysis was performed comparing WC and AT mice, and 896 up-regulated and 908 down-regulated genes were detected with FDR=5% and absolute fold change greater than 2 (Figure 6B). Figure 6B shows the expression patterns of these differentially expressed genes (DEGs) where each row represents a gene and each column represents a sample. Ingenuity pathway analysis was performed on these DEGs and top pathways are shown in Supplemental Figure S3. In accord with the results described above, the Wnt/beta-catenin and NRF2-mediated oxidative stress response pathways were highly enriched, with substantial numbers of responsive pathway components (Figure 6E, F).

**Figure 6.**
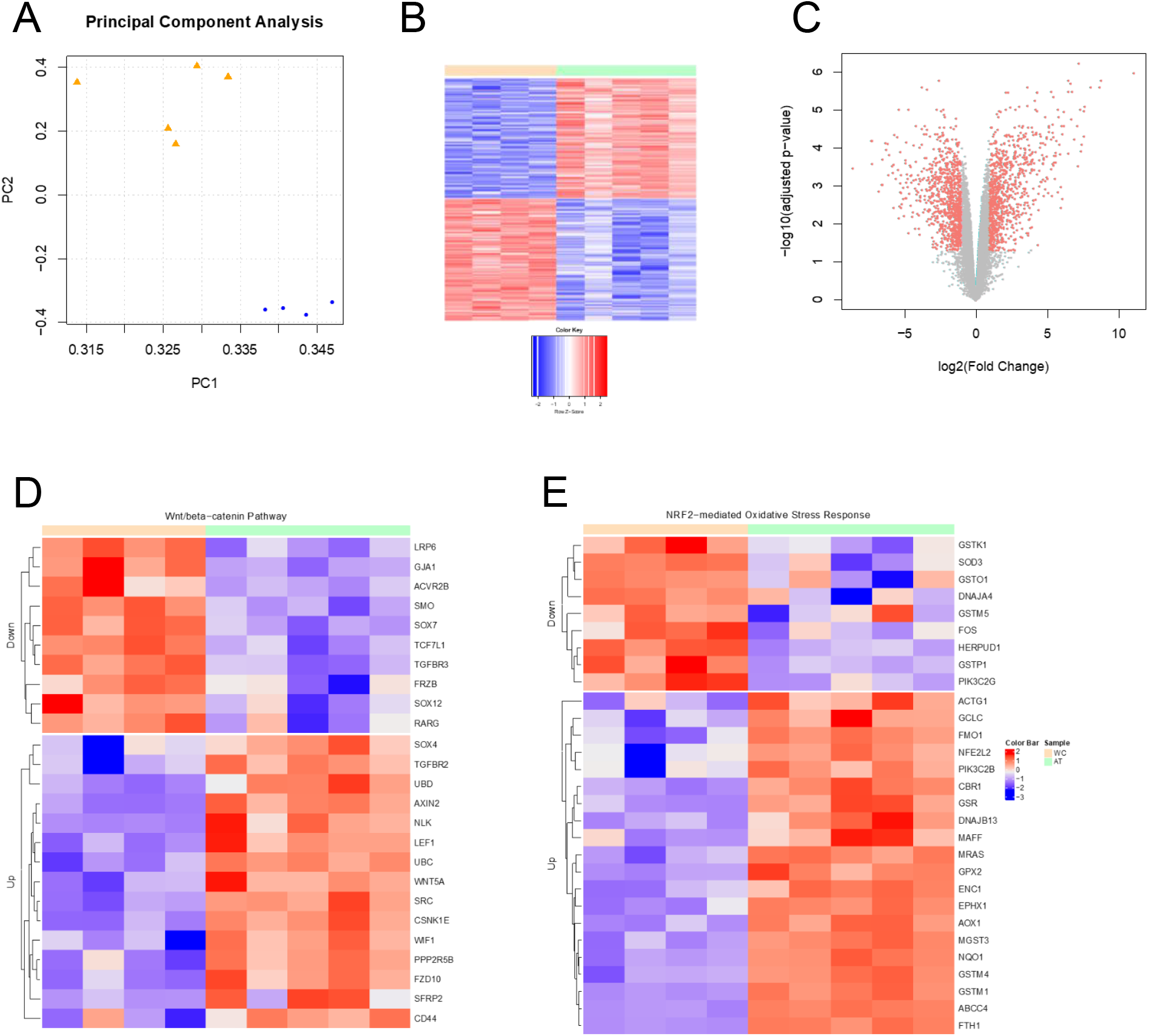
Transcriptome analysis on mouse model. (A) Principal component analysis on the 9 mouse samples. Four wild type samples are marked as blue dots and five AT samples are marked as orange triangles. (B) Volcano plot for differential expression analysis, with x-axis for log2 fold change and y-axis for −log10(adjusted p-values). (C) Heatmap on differentially expressed genes, with rows for genes and columns for samples. (D) Differentially expressed genes in Wnt/beta-catenin pathway. (E) Differentially expressed genes in NRF2-mediated oxidative stress response pathway.

### CAR + HBx mouse tumors share conserved pathways and gene signatures with human HBV-associated HCC tumors

HBx is a well-known factor involved in HBV-positive HCC cases, but the role of CAR in human HCC is less well defined. We previously showed a conserved gene signature between a mouse model using CAR activation and β-catenin mutated human HCC (15). Additionally, CAR has been identified as a key gene responsible for mediating the progression of non-alcoholic fatty liver disease (NAFLD) to HCC, suggesting that CAR may be more highly activated in early-stage tumors (34). To test whether our mouse-derived tumors resembled human HBx tumors, we analyzed three data sets containing large cohorts of HBV-positive patients: GSE94660, GSE121248, and TCGA. Using these data sets, we performed differential gene expression and pathway analysis, following the pipeline used for analysis of our mouse model.

In order to compare the similarity between mouse model and human study, we first performed differential expression and pathway analysis on mouse and human studies individually, and then explored their molecular mechanisms in two comparisons (Supplemental Figure S4). First, mouse gene signatures were validated in human studies. Mouse DEGs (Figure 6) were first mapped to homologous human genes by Mouse Genome Informatics (MGI) and then compared with human DEGs. Gene expression analysis showed a shared set of genes between mice and each human data set (Figure 7A-C).

**Figure 7.**
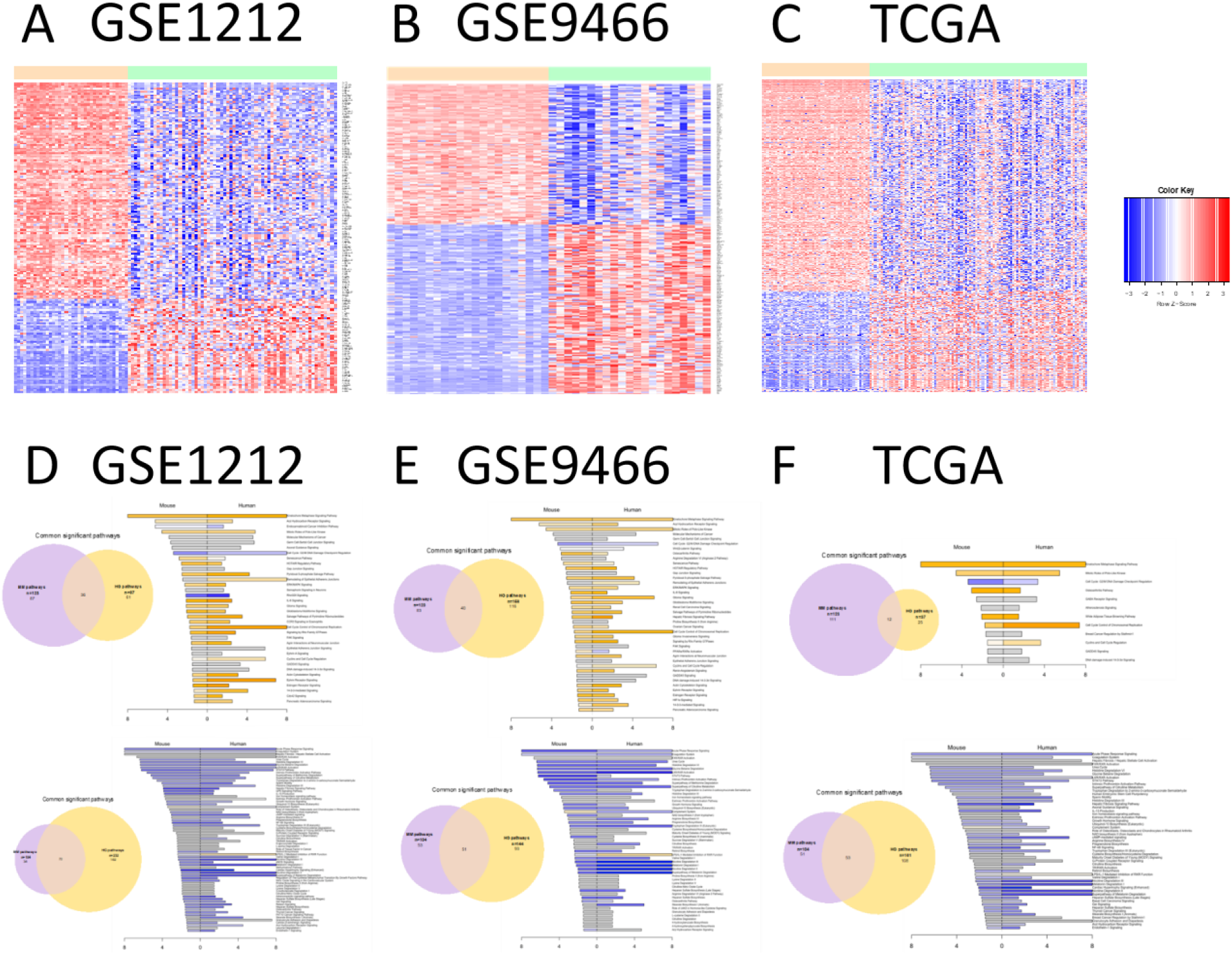
Common gene signatures and pathways conserved in HBV-positive HCC tumors and ATX + TCPOBOP mice. (A-C) Gene signatures of three human data sets mapped to differentially expressed genes within the mouse tumor. (D-F) Top conserved pathways between mouse and human.

To identify common differentially regulated pathways, we performed IPA analysis on top up- or downregulated genes. In the analysis of mouse tissue, we detected 123 significant pathways from up-regulated genes; FDR=5% (Figure 7D). Similarly, we identified 97 significant pathways in the human GSE121248 data set. When we checked their similarity, 36 common pathways were found between the mouse and human data sets. That is, 30% of the pathways that altered in our mouse model are similarly changed in human HBV patients. Similar analysis was performed on the down-regulated genes, where 70 common pathways (out of 104 mouse pathways) were identified between mouse and human GSE121248 (Figure 7D). For the other two human studies, very consistent and robust results were drawn on the pathway similarity. Additionally, on the top of the blindly searching of enriched pathways, we specifically checked Wnt/beta-catenin pathway (Figure 8A-C) and NRF2-mediated oxidative stress response pathway (Figure 8D-F). The alterations of both pathways were validated in all the three human studies. Pathway analysis showed conserved pathways between each of the data sets and our mouse tumors (Figure 7D-F). Among the top conserved pathways within each data set were kinetochore metaphase signaling pathways, hepatic fibrosis pathways, and G2/M DNA damage checkpoint regulation pathways.

**Figure 8.**
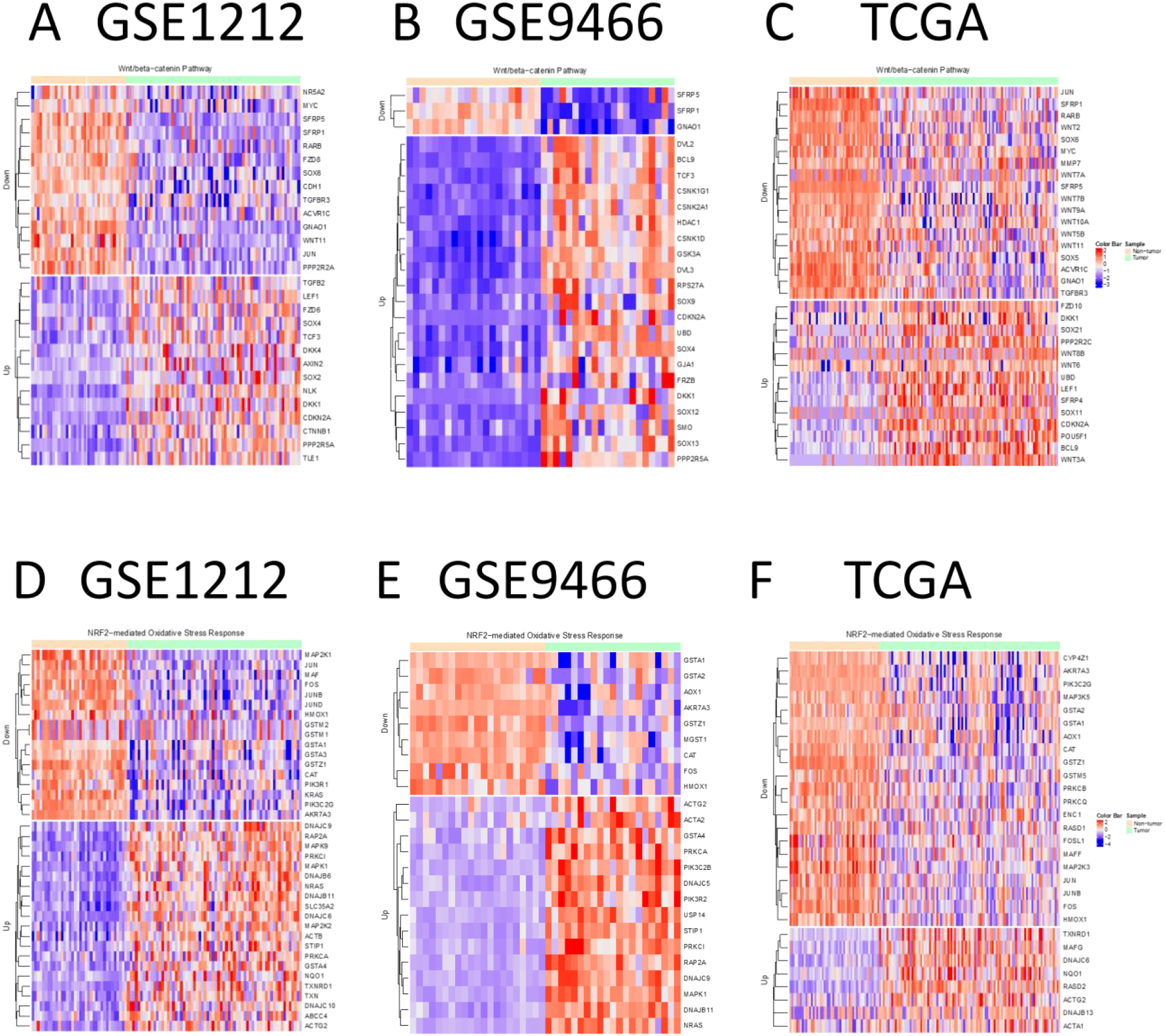
Differentially expressed genes in Wnt/beta-catenin pathway and NRF2-mediated oxidative stress response pathway.

In addition to investigating the relationship of our tumor model with HBV-associated HCCs, we wanted to determine whether our model more closely resembles HBV-associated HCCs compared with non-HBV-associated HCCs. We therefore applied a hierarchical clustering analysis to TCGA data containing HBV, non-HBV, and normal liver tissue samples. Using the top 594 differentially expressed genes from our mouse model, we found that although normal tissue forms its own cluster, there was no separation of HBV- and non-HBV-associated tumors (Supplemental Figure S5). This result was not surprising, as previous clustering analysis has indicated that the two etiologies do not divide into distinct groups (35). Therefore, it is likely that our model can be applied equally well to both HBV- and non-HBV-associated HCC tumors.

## Discussion

In this study, we demonstrate a clear functional synergy between the nuclear receptor CAR and the HBV X protein in hepatocarcinogenesis. HBx, the main regulatory protein encoded by HBV, is not considered to be directly transforming but plays a significant positive role in HBV-associated hepatocarcinogenesis (10). HBx has been reported to support β-catenin activation, which is highly associated with HBV-positive HCC (26,36,37). However, transgenic mice expressing HBx in hepatocytes under control of the α-1-antitrypsin promoter (ATX mice) do not develop tumors at higher rates than wild-type controls (21), and we did not observe significant β-catenin activation in 12-month-old ATX mice without other intervention. β-catenin activation is also highly promoted in HCC mouse models that utilize CAR activation as a tumor promoter (14,38), suggesting a possible functional synergy between HBx and CAR. Here we have demonstrated that CAR activation in mice harboring the HBx transgene results in a significant tumor burden, and that the tumors are characterized by widespread β-catenin activation. Functional interaction between CAR and β-catenin has been well-documented. Approximately 80% of phenobarbital-mediated mouse liver tumors contain activating mutations in β-catenin (14). Our lab previously showed that direct, simultaneous activation of both CAR and β-catenin leads to bypass of senescence checkpoints and stimulates both rapid and persistent liver proliferation in the short term, and tumorigenesis in the longer term (15). There are multiple explanations for the apparent selection of β-catenin activation in CAR-mediated tumor promotion models. CAR activation, in addition to conferring a selective advantage to cells with carcinogen-induced β-catenin activation (14), may also induce β-catenin mutations on its own (39). CAR has been shown to interact with the Wnt/β-catenin pathway and induce β-catenin nuclear localization through Akt signaling and may also crosstalk with β-catenin through the Hippo/YAP signaling pathway (40,41). In the current study, we found that HBx expression on its own was associated with modest β-catenin activation at early timepoints, but that activation dropped off in older mice. In contrast, tumors arising in TCPOBOP-treated HBx transgenic mice have substantial β-catenin activation. This suggests that CAR contributes to the selective expansion of cells with dysregulated β-catenin pathways, though it does not rule out the possibility that CAR, along with HBx, directly contributes to β-catenin nuclear localization. Surprisingly, CAR appears to inhibit β-catenin target gene expression at early timepoints. This unexpected result suggests a transient inhibitory crosstalk between CAR and β-catenin, a previously undocumented finding that warrants further study.

FoxM1 is a key downstream proliferative effector in the CAR-activated liver. When activated, CAR binds the c-Myc promoter, which in turn transcriptionally activates FoxM1, a transcription factor that transactivates expression of cell cycle control genes (42). FoxM1 reportedly also interacts with the Wnt/β-catenin pathway to promote nuclear translocation of β-catenin (28). In our study, we found both early activation of FoxM1 in TCPOBOP-treated mice and sustained FoxM1 activation in tumors. This raises the possibility that CAR may play an indirect role in β-catenin activation through the induction of FoxM1. FoxM1 and CAR are also each involved in promoting Nrf2 activation (29,31). Nrf2 mediates the antioxidant response and has been documented as an early driver of HCC (43). Although Nrf2 has a protective role in healthy liver, sustained activation of Nrf2 is associated with more aggressive HCC and lower overall patient survival (44). In this study, we have documented both increased β-catenin activity and increased expression of Nrf2 targets, two events common in human HCC. We suggest that these features in the mouse CAR/HBx tumors may recapitulate those of the subset of human HCC tumors that feature both β-catenin and Nrf2 hyperactivation.

Interestingly, β-catenin mutations are less likely to occur in HBV-positive HCC than in other HCC etiologies (45,46), yet there is a positive correlation between HBx expression in human HCC samples and nuclear localization of β-catenin (47). Over the course of our study, we explored potential SNVs/mutations correlated with our mouse tumor model. We first performed SNV/mutation calling on each individual sample, and then annotated the SNVs positions to genes. Previous studies have identified gene mutations that are highly correlated with HCC (48). We specifically targeted these genes and counted the number of SNVs across the WC and AT samples (Supplemental Figure S6). From them, TSC2, SYNE2, RB1, WNT2, FRAS1 and NFE2L2 showed the most SNVs, while CTNNB1 mutations appeared to be relatively rare. We further narrowed all the SNVs/mutations down to 19 SNVs that weren’t detected in any WC samples but were observed in at least 3 AT samples. These SNVs can be explored further for functional analysis, including direct test of DNA alterations on paired adjacent tumor and tumor samples are generally used to detect somatic mutation.

HCC is refractory to traditional chemotherapy treatment, and until recently, the only approved treatments for intermediate to advanced stage HCC were multi-kinase inhibitors sorafenib and lenvatinib, which extend life expectancy by only a few months (49). Unlike other tumors, HCC does not have clearly defined functional subtypes, but there are functional differences based on genetic etiologies and other factors (TCGA Cell). Therefore, it is imperative to identify targeted intervention strategies based on tumor subtype, and by extension, to identify appropriate models for HCC subtypes. In order to study specific types of HCC, we need models that align well with various tumor subtypes. Additionally, some questions have been raised about whether CAR activation in humans causes direct DNA replication, and by extension, whether CAR is relevant to human HCC (50). Although the answers to these questions are still being investigated, our model shows a conserved gene signature with three independent sets of HBV-related HCC. We additionally identified strong conservation of several key pathways involved in both mouse and human cancers. This indicates that CAR-initiated mouse tumors are related to human HCC, regardless of initiation mechanism.

Our results reveal a novel interplay between CAR activation and expression of HBx. While HBx causes transient, mild upregulation of β-catenin transcriptional targets, we found that this effect was not present in older mice. The addition of CAR activation, however, apparently selects for β-catenin activation in tumors. Intriguingly, CAR activation in the short term had the opposite effect, serving to suppress β-catenin transcriptional targets. This supports the hypothesis that CAR activation creates an environment supportive of the clonal expansion of β-catenin activated cells rather than directly activating the β-catenin pathway. Our study indicates that upregulation of FoxM1 and Nrf2 are key events in tumorigenesis in this model. Additionally, consistent with human HBV-mediated HCC, *CTNNB1* mutations were scarce in our tumors despite strong β-catenin nuclear localization. Our results demonstrate a conserved gene signature with three independent sets of HBV-positive human HCC tumors, as well as sharing conserved pathways with these data sets. Together, these data identify the relevance of our model for modeling human HCC.

