## Supplemental File for "Constitutive Androstane Receptor and Hepatitis B Virus X Protein Cooperatively Induce β-catenin-Activated Liver Tumors"

**Supplemental Figure S1. Results of ATX genotyping.** Presence of ATX transgene was confirmed by genotyping. A clear band at 500 kB is amplified in mice that are positive for the transgene. No band is amplified in wild-type mice. HBx RNA expression was confirmed by RT-qPCR. ** $p < 0.01$. Data are represented as mean ± SEM.

**Supplemental Figure S2. Tissue staining in 10 month CAR -/- samples.**

**Supplemental Figure S3.** Top significant pathways on mouse model.

**Supplemental Figure S4.** Comparison between mouse model and human studies. (Pipeline) Analysis pipeline to compare mouse and human studies.

**Supplemental Figure S5.** Hierarchical clustering on human TCGA samples employing the gene signatures selected from the mouse model.

**Supplemental Figure S6.** SNVs calling on mouse RNA sequencing data. (A) Heatmap for number of SNVs called on HCC-related genes across all the samples. (B) Heatmap for SNV rate on the 19 selected positions across all the samples.

**Figure S1**

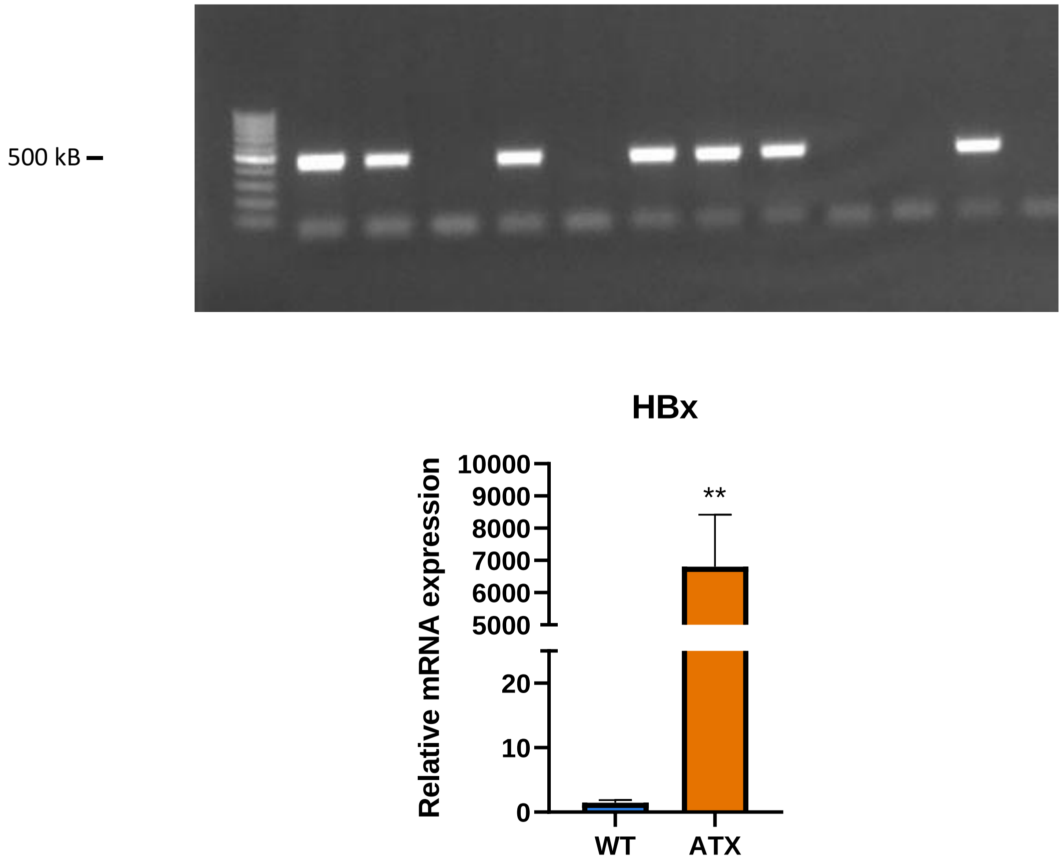
500 kB
HBx
Relative mRNA expression
10000
9000
8000
7000
6000
5000
20
10
0
**
WT
ATX

**Figure S2**

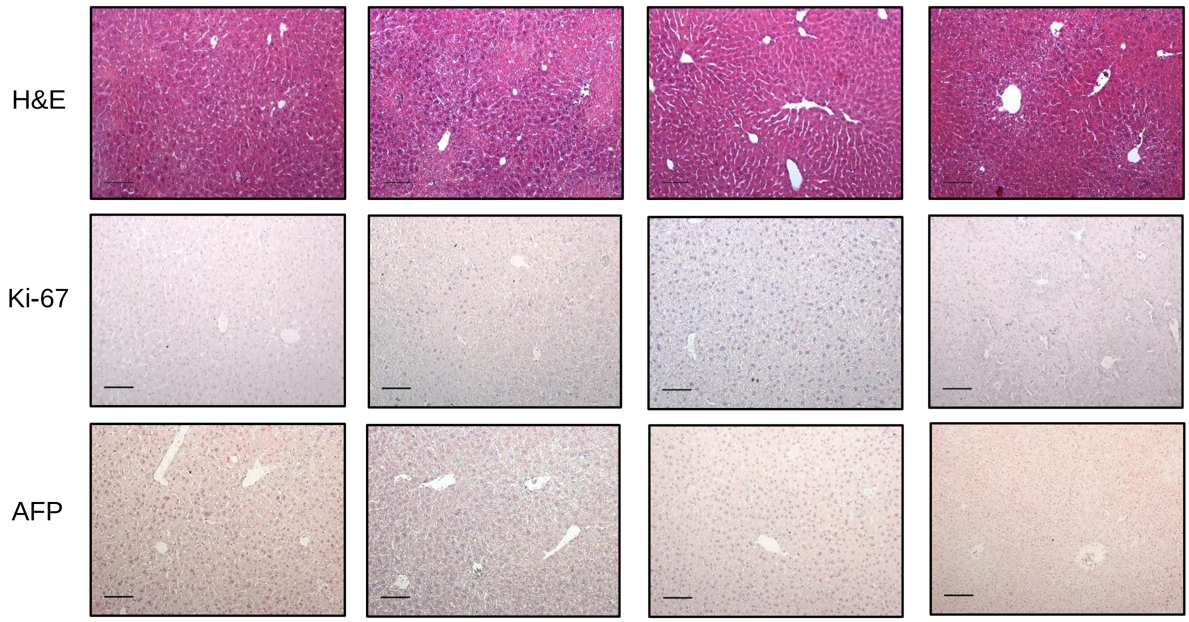

### Figure S3

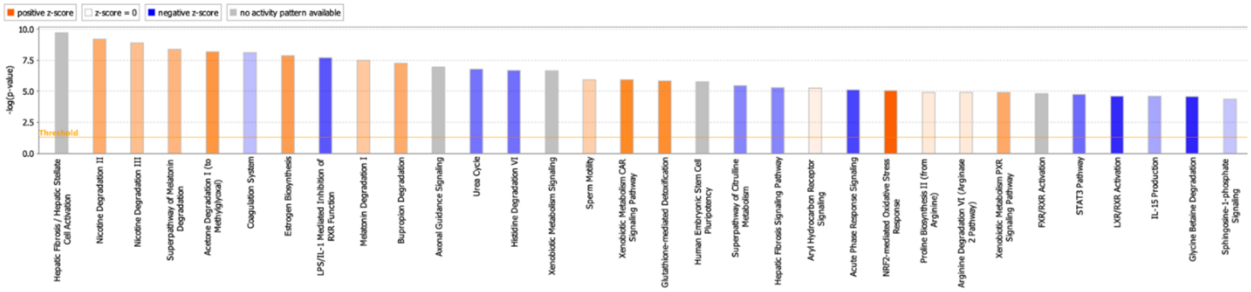

positive z-score
z-score = 0
negative z-score
no activity pattern available
-log(p-value)
10.0
7.5
5.0
2.5
0.0
Threshold
Hepatic Fibrosis / Hepatic Stellate Cell Activation
Nicotine Degradation II
Nicotine Degradation III
Superpathway of Melatonin Degradation
Acetone Degradation I (to Methylglyoxal)
Coagulation System
Estrogen Biosynthesis
LPS/IL-1 Mediated Inhibition of RXR Function
Melatonin Degradation I
Bupropion Degradation
Axonal Guidance Signaling
Urea Cycle
Histidine Degradation VI
Xenobiotic Metabolism Signaling
Sperm Motility
Xenobiotic Metabolism CAR Signaling Pathway
Glutathione-mediated Detoxification
Human Embryonic Stem Cell Pluripotency
Superpathway of Citrulline Metabolism
Hepatic Fibrosis Signaling Pathway
Aryl Hydrocarbon Receptor Signaling
Acute Phase Response Signaling
NRF2-mediated Oxidative Stress Response
Proline Biosynthesis II (from Arginine)
Arginine Degradation VI (Arginase 2 Pathway)
Xenobiotic Metabolism PXR Signaling Pathway
FXR/RXR Activation
STAT3 Pathway
LXR/RXR Activation
IL-15 Production
Glycine Betaine Degradation
Sphingosine-1-phosphate Signaling

**Figure S4**

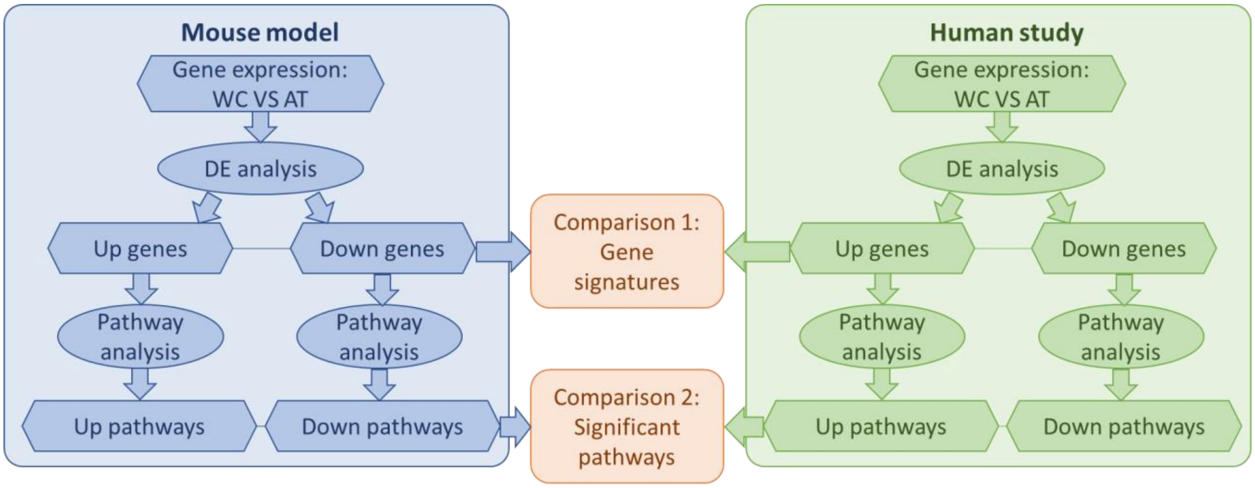
Mouse model
Gene expression: WC VS AT
DE analysis
Up genes
Down genes
Pathway analysis
Pathway analysis
Up pathways
Down pathways
Comparison 1: Gene signatures
Comparison 2: Significant pathways
Human study
Gene expression: WC VS AT
DE analysis
Up genes
Down genes
Pathway analysis
Pathway analysis
Up pathways
Down pathways

**Figure S5**

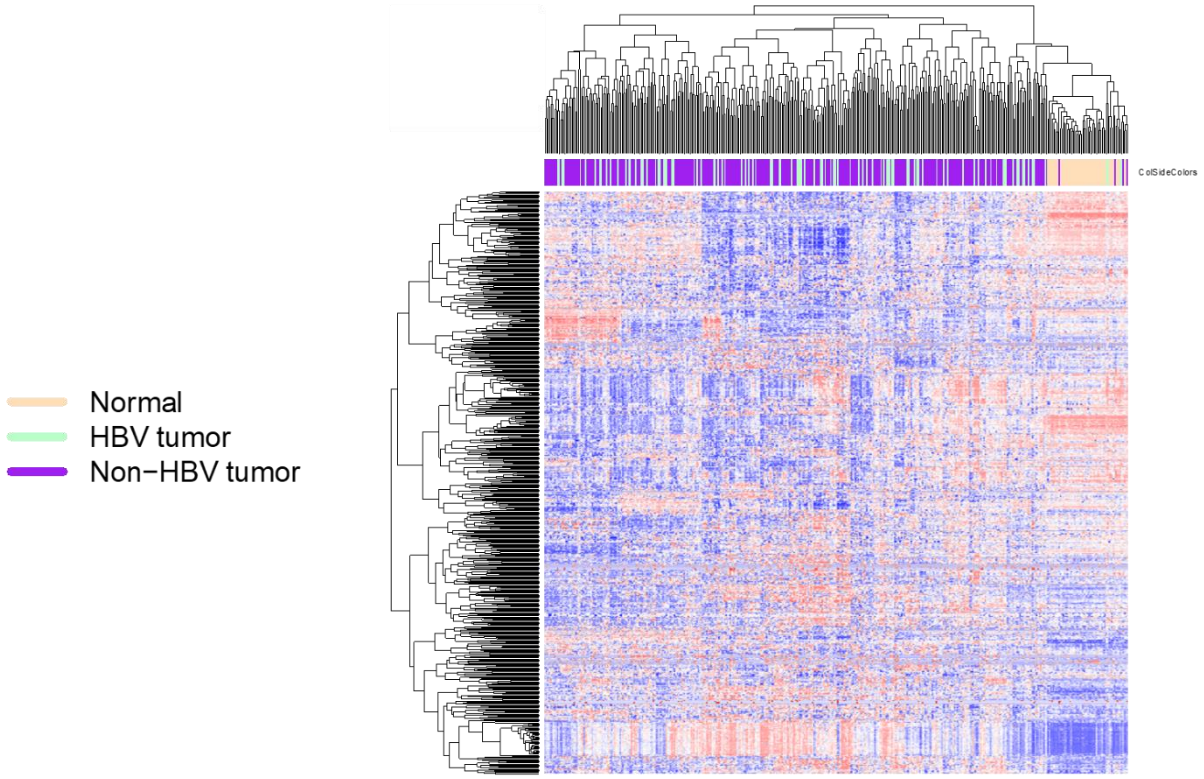

**Figure S6**

A

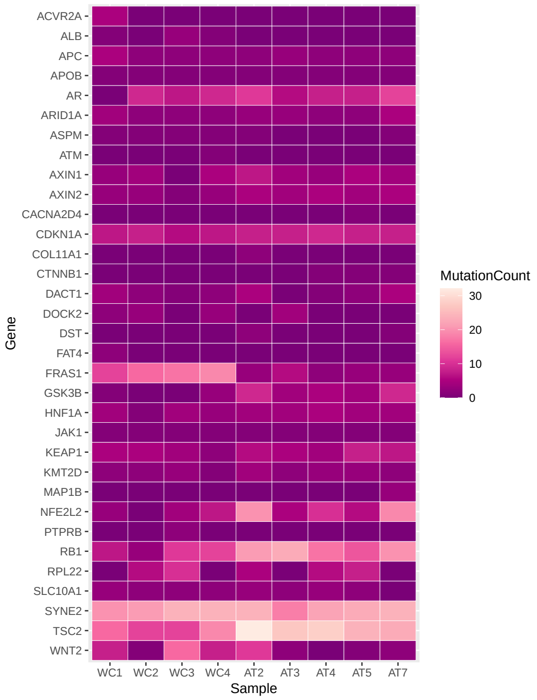
ACVR2A
ALB
APC
APOB
AR
ARID1A
ASPM
ATM
AXIN1
AXIN2
CACNA2D4
CDKN1A
COL11A1
CTNNB1
DACT1
DOCK2
DST
FAT4
FRAS1
GSK3B
HNF1A
JAK1
KEAP1
KMT2D
MAP1B
NFE2L2
PTPRB
RB1
RPL22
SLC10A1
SYNE2
TSC2
WNT2
Gene
WC1 WC2 WC3 WC4 AT2 AT3 AT4 AT5 AT7
Sample
MutationCount
30
20
10
0

B

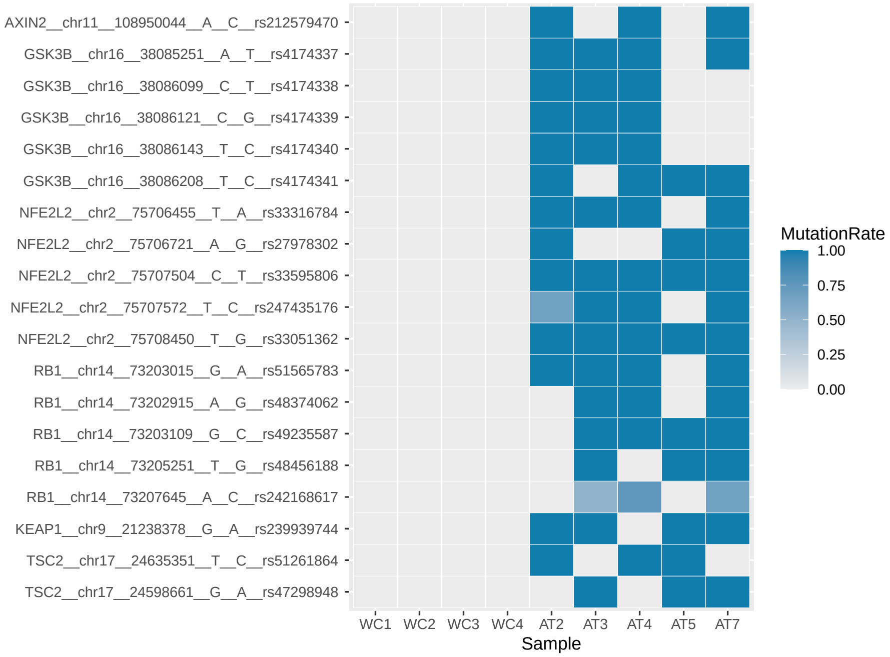
AXIN2__chr11__108950044__A__C__rs212579470
GSK3B__chr16__38085251__A__T__rs4174337
GSK3B__chr16__38086099__C__T__rs4174338
GSK3B__chr16__38086121__C__G__rs4174339
GSK3B__chr16__38086143__T__C__rs4174340
GSK3B__chr16__38086208__T__C__rs4174341
NFE2L2__chr2__75706455__T__A__rs33316784
NFE2L2__chr2__75706721__A__G__rs27978302
NFE2L2__chr2__75707504__C__T__rs33595806
NFE2L2__chr2__75707572__T__C__rs247435176
NFE2L2__chr2__75708450__T__G__rs33051362
RB1__chr14__73203015__G__A__rs51565783
RB1__chr14__73202915__A__G__rs48374062
RB1__chr14__73203109__G__C__rs49235587
RB1__chr14__73205251__T__G__rs48456188
RB1__chr14__73207645__A__C__rs242168617
KEAP1__chr9__21238378__G__A__rs239939744
TSC2__chr17__24635351__T__C__rs51261864
TSC2__chr17__24598661__G__A__rs47298948
WC1 WC2 WC3 WC4 AT2 AT3 AT4 AT5 AT7
Sample
MutationRate
1.00
0.75
0.50
0.25
0.00

**Table S1. Primer sequences for RT-qPCR and genotyping primers.**

#### Table S1

| Primer | Forward Sequence | Reverse Sequence |
|---|---|---|
| AFP | CATGCTGCAAAGCTGACAA | CTTTGCAATGGATGCTCTCTT |
| ATX | ATGGCTGCTAGGCTGTACTG | GTACAAGAGATGATTAGGCAG |
| CAR | GCATCCTACACCCGATCTTG | AGTTCCTCGGCCCATATTCT |
| CCNB2 | GCCAAGAGCCATGTGACTATC | CAGAGCTGGTACTTTGGTGTTC |
| Cyclophilin A | CAAGACTGAATGGCTGGATG | ATGGGTAAAATGCCCGC |
| Cyp2B10 | AAGCTCATTCTCCAGCCAGA | CTGTGGGCACCAGGAAAG |
| FoxM1 | ACTTTAAGCACATTGCCAAGC | GGAGAGAAAGGTTGTGACGAA |
| FoxO1 | ATGCTCAATCCAGAGGGAGG | ACTCGCAGGCCACTTAGAAAA |
| GPR49 | GGACCAGATGCGATACCGC | CAGAGGCGATGTAGGAGACTG |
| GS | CTGAGTGGAACTTTGATGGCT | GGAAGGGGTCTCGAAACATGG |
| Gstp1 | TGTCACCCTCATCTACACCAAC | GGACAGCAGGGTCTCAAAAG |
| NFE2l2 | CATGATGGACTTGGAGTTGC | CCTCCAAAGGATGTCAATCAA |
| Nqo1 | AGCGTTCGGTATTACGATCC | AGTACAATCAGGGCTCTTCTCG |
| Slc1a2 | GCCAACAATATGCCCAAGCAG | GACACCAAACACAGTCAGTGA |
| Sqstm1 | AGACCCCTCACAGGAAGGAC | CATCTGGGAGAGGGACTCAA |
| TCF | AGCTTTCTCCACTCTACGAACA | AATCCAGAGAGATCGGGGGTC |
| TGF-b | CTGGCGAGCCTTAGTTTGGAC | CCACCTGCAAGACCATCGAC |
